# A combination of the extract from *Phellinus linteus* with *Smilax* spp. attenuates colorectal and prostate cancers

**DOI:** 10.1101/2025.10.19.683316

**Authors:** Chiratchaya Chongrak, Peerapat Visitchanakun, Prakaithip Somjit, Panomwat Amornphimoltham, Asada Leelahavanichkul, Pattamawadee Yanatatsaneejit

## Abstract

The current study investigated the effects of Thai remedy herbal extracts from Phellinus linteus (PL) and a combination of P. linteus with Smilax corbularia and S. glabra (PSS) on colorectal, prostate cancer cell lines, along with kidney cell lines *in vitro*, as well as their effects on mice subcutaneously injected with colorectal cancer cells. The IC_50_ values of PSS and PL were evaluated using the MTT assay, and their effects on cell cycle distribution and apoptosis induction were examined. RT-qPCR was performed to analyze the expression of proliferation-related genes following treatment with PSS and PL. Tumor-associated macrophages (TAMs) were also treated with these extracts to assess gene expression related to macrophage polarization, and macrophages’ tumoricidal activity was measured using CFDA-SE–labeled cancer cells. In addition, an animal tumor xenograft model was employed to determine the in vivo efficacy of PSS and PL. The results revealed that both PSS and PL exerted cytotoxic effects on colorectal, prostate, and kidney cancer cell lines. PSS induced G2 phase arrest in all colorectal cancer cell lines but not in PC3 cells and showed a mild effect on apoptosis induction in colorectal and prostate cancer cells. The inhibition of proliferation was partly associated with the downregulation of genes involved in cancer progression by both PL and PSS. Moreover, PL and PSS interfered with cancer supernatant–stimulated macrophages, upregulating M1-associated genes and downregulating M2-associated genes. In vivo, PSS reduced tumor volume in HCT116 cell–injected mice, and co-incubation of fluorescently labeled HCT116 cells with TAMs and PSS decreased fluorescence intensity, whereas PL alone did not exhibit these effects. Collectively, these findings indicate that both PL and PSS inhibit cancer cell proliferation *in vitro*, but only PSS demonstrated significant antitumor activity *in vivo*.

## Introduction

Cancer is a leading cause of death worldwide, especially colorectal and prostate cancer (1), and there are a spectrum of interventions, including surgery, chemotherapy, hormonal treatment and targeted therapies, which are tailored to the stage and type of cancer (2). Since mortality rates and the treatment side effects of cancer are still high, several natural compounds have gained attention in the adjuvant therapy of cancer, with the aim of improving survival and reducing the side effects of the standard treatments (3). Herbal therapy is becoming increasingly popular as a complement to conventional cancer treatments. The information for the proposed approach is still preliminary, but overall, it is promising (4). Various phytochemicals are derived from medicinal plants such as curcumin, ellagic acid, EGCG, berberine, artemisinins, ginseng, gallic acid artesunate and beta-glucan, have been shown to have anticancer properties, including antiproliferative, antiangiogenic, proapoptotic, and antimetastatic effects. They also control autophagy, reverse drug resistance, improve immunity, and aid in chemotherapy development in both vitro and in vivo studies (5).

In Thailand, several herbs have traditionally been used in cancer treatment; for example, *Smilax* spp., especially *Smilax corbularia* (SC) and *S. glabra* (SG) (also referred to as catbriers, greenbriers, prickly ivy and smilaxes), are climbing flowering plants in tropical and subtropical regions that possess anticancer properties, mostly from phenolic compounds (6–9). Additionally, *Smilax* spp. are well-known herbs for the treatment of rheumatism, syphilis, diabetes mellitus and cancer (9–11). On the other hand, *Phellinus linteus* (PL), a medicinal mushroom, is well-known in alternative medicine due to its anticancer effects, partly through the inhibition of cell proliferation and the induction of apoptosis of malignant cells (12, 13). Accordingly, (1→3)/(1→6)-β-glucan, the major cell wall component of mushrooms, has been demonstrated to exert an antitumor effect on several animal models (14, 15), possibly through the immune modulation of glucan on numerous immune cells (16, 17). While tumor-associated macrophages (TAMs) facilitate the proliferation of malignant cells (18), the administration of glucan reduces the beneficial effects of TAMs on cancer and enhances the tumoricidal activities of TAMs, partly through an alteration in cell energy status (14). Notably, the anticancer effect of herbal medicine might not only have a direct impact on the proliferation of malignant cells, but also on immune activation. Nevertheless, the *in vitro* assessment of herbal extracts on cancer mostly relies on the co-incubation of the substances with malignant cell lines (14, 19–22), which might not cover all possible mechanisms. *In vitro* experiments on immune cells, as well as *in vivo* studies, are necessary to determine the translational use of herbal extracts in cancer therapy.

Since *Phellinus*, an edible mushroom, is commonly found in China, Japan, South Korea and Thailand, and *Smilax* spp. plants are distributed throughout the eastern and south-eastern regions of Asia, the mixtures of PL, SC, and SG (denoted as PSS) appear in several traditional Thai medicines. Recently, we demonstrated the direct anti-proliferative activity of PSS on several breast cancer cell lines, which might be useful as an additional treatment (12). Despite the obvious molecular differences between breast cancer, colorectal and prostate cancer, the effect of PSS on cancer, at least in part, might not be specific to the cancer cell type. Due to the high incidence of colon and prostate cancer cases in the Thai population (23, 24), PSS has been proposed as a potential herbal therapy for colon and prostate cancer in some Thai patients with cancer. Given the potential advantages of PSS against these tumors, as well as the absence of *in vitro* and *in vivo* studies for PL and PSS in these cancer cell types, the present study investigated the impact of PSS in both *in vitro* and *in vivo* experiments. We also addressed the mechanisms involved in cytotoxicity against these cancer cells, such as anti-proliferation through cell cycle arrest, apoptosis and gene-related cell proliferation. In addition, it was hypothesized that the combination of PL and PSS may be effective in the attenuation of malignancy, partly through an impact on TAMs.

## Material and methods

### Cell culture

Two colon cancer cell lines (HCT116 and SW620), one colorectal cancer cell line (HT-29), one prostate cancer cell line (PC3) and one kidney cell line (HEK 293) were purchased from the American Type Culture Collection (ATCC, VA, USA), and short tandem repeat profiling was performed for authentication. These cell lines were cultured in Gibco^®^ Dulbecco’s modified Eagle’s medium (DMEM; Thermo Fisher Scientific, MA, USA) supplemented with 10% Gibco fetal bovine serum (FBS; Thermo Fisher Scientific) and 1% Gibco antibiotic-antimycotic (Thermo Fisher Scientific), and were maintained in a humidified atmosphere with 5% CO_2_ at 37°C.

### Preparation of PL and PSS extracts

All *in vitro* procedures were carried out as previously reported (10). In brief, for PSS, crude powders from the fruiting body of PL, and the rhizomes of SC and SG (M herbs inter Ltd., Bangkok, Thailand) at a ratio of 3:1:1 (PL:SC:SG) were extracted using distilled water with a solid-to-liquid ratio of 1:50 in a Soxhlet apparatus at 80°C for 5 h, after which they were filtered (Whatman No. 1 paper) and oven-dried at 70°C overnight. The ratio of PL:SC:SG in the PSS combination employed in the present study was based on a traditional Thai recipe, and the concentrations of PL and PSS were determined by a recent publication indicating the advantages of both drugs in breast cancer (10). The extraction yield was calculated using the following formula: Yield (g/g) = extract obtained after evaporation (g)/dry weight (g).

### MTT assay

The MTT method was used to assess the cytotoxicity of the extracted samples against each cell line. In brief, all cell lines (5×10^3^ cells) were plated in 96-well plates and incubated at 37°C for 24 h. The cells were then treated with PSS and PL at various dosages (75-650 µg/ml and 250-3,750 µg/ml, respectively) for 48 h in colon cancer cell lines and a kidney cell line, and for 72 h in the prostate cancer cell line. Untreated cells were used as a negative control. Cisplatin (Sigma-Aldrich) was supplied as a positive control at doses ranging between 2.12 and 4.08 µg/ml. After 48 h or 72 h of treatment, the MTT assay was employed to determine the half-maximal inhibitory concentration (IC_50_). Briefly, after removing the extract medium, each well was incubated with 5 mg/ml MTT dissolved in fresh medium at 37°C for 3 h. After discarding the solution, DMSO was added and incubated at 27°C for 5 min. Absorbance was determined using a spectrophotometer at 492 nm and a reference wavelength of 630 nm for all samples. The inhibition percentages for untreated, PSS-, PL- and cisplatin-treated groups were compared, and presented as percentages of the untreated control. The results of the MTT assay (IC_50_-dependent) were applied to conduct further cell proliferation and apoptosis assays.

### Cell cycle analysis

Flow cytometric analysis with propidium iodide (PI; DNA staining) was performed using colon and prostate cancer cells (5×10^5^ cells/well) treated with PSS, PL and cisplatin (IC_50_ values from the MTT experiment) for 48 and 72 h, respectively. Cells were trypsinized, fixed with cold 70% ethanol at 4°C overnight and centrifuged at 14,000 rpm for 15 min, after which, 50 µl of a 100 µg/ml stock of RNase (Sigma-Aldrich) was added and the cells were stained with PI solution (BD Pharmingen, CA, USA) before analysis using a flow cytometer. Cell cycle arrest at G_0_/G_1_, S and G_2_ phases was evaluated.

### Apoptosis assay

The Annexin V-DY-634/PI apoptosis detection kit (Abcam, Cambridge, UK) was utilized to assess apoptosis. After seeding and treating the cells with the IC_50_ values of PSS, PL and cisplatin as aforementioned, the cells were collected and washed twice with PBS. The cells were then stained with DY-634 and PI, as per the manufacturer’s instructions. Then, a flow cytometric analysis was performed. Early and late apoptosis, and necrosis were determined by the low and high levels of Annexin V and PI; high levels of Annexin V and low levels of PI indicated early apoptosis, high levels of both Annexin V and PI indicated late apoptosis, and low levels of Annexin V and high levels of PI indicated necrosis.

### Gene expression analysis

The colon, colorectal and prostate cancer cells that had been treated for 48 or 72 h were trypsinized, and RNA extraction was performed using Invitrogen^®^ TRIzol™ (Thermo Fisher Scientific). Subsequently, 750-1,000 ng total RNA from each sample was used to synthesize cDNA using a cDNA synthesis kit (Biotechrabbit, Hennigsdorf, Germany), as instructed. The cDNA underwent reverse transcription-quantitative PCR (RT-qPCR) using 4X CAPITAL™ qPCR Green Master Mix (Biotechrabbit) with forward and reverse candidate gene primers, as shown in Table I. *GAPDH* served as an internal control (25–28). qPCR amplification was performed using a QuanStudio™ 5 Real-time PCR System (Thermo Fisher Scientific) with the following thermocycling conditions: 40 cycles at 95°C for 15 sec, 60°C for 30 sec and 72°C for 45 sec. The 2^-ΔΔCq^ technique (29) was used to compare gene expression changes between mixture- or PL extract-treated and untreated cells.

**Table 1.**
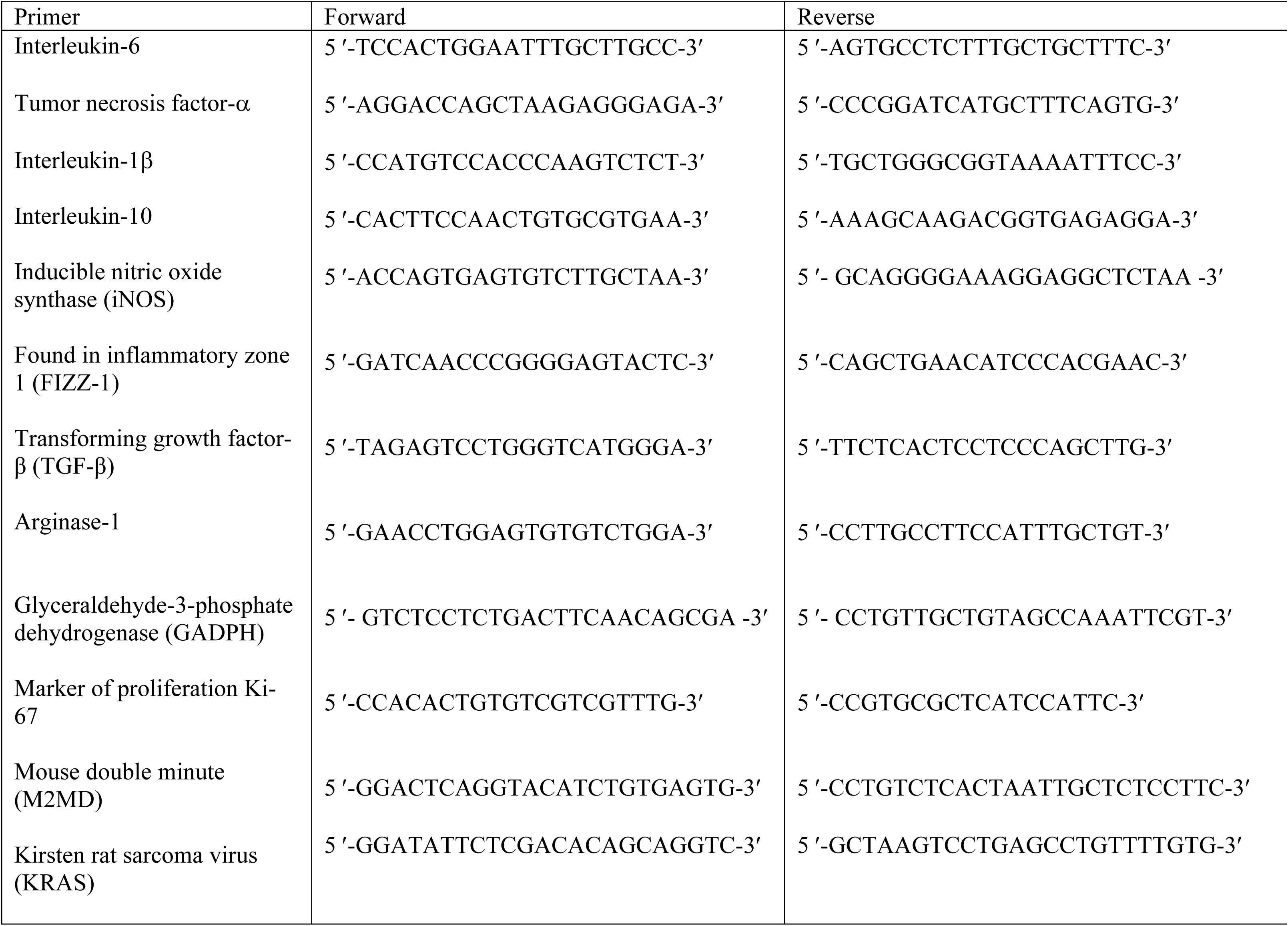
List of primers.

### Macrophage experiments

Macrophages were derived from human monocytoid THP-1 cells (TIB-202; ATCC) in Roswell Park Memorial Institute medium (RPMI; Thermo Fisher Scientific) supplemented with 10% FBS, 1% Gibco antibiotic-antimycotic and 50 ng/ml phorbol 12-myristate 13-acetate (Sigma-Aldrich). The induction of TAMs was performed as previously described (14, 30, 31). Because of the effects of secreted chemicals from cancer cells on macrophages, such as succinate, osteopontin and other metabolic byproducts, macrophages recruited to cancer microenvironments are transformed into TAMs (18). As a result, the use of tumor-conditioned medium containing a number of excreted compounds to convert macrophages into TAMs *in vitro* is well established (14, 32). To prepare the conditioned medium for cancer cells (HCT116), 3×10^6^ cells were incubated in 9 ml modified DMEM for 72 h, centrifuged at 1,500 rpm for 10 min at 4°C and filtered through 0.22-μm filters. Macrophages (1×10^6^ cells/well) were treated with 500 µl/well HCT116 conditioned medium for 24 h before being removed with a pipette, rinsed with PBS and cultured in standard DMEM.

TAMs were incorporated into extracts (PL or PSS) or DMEM (control) for 24 h. Cytokines (TNF-α, IL-6 and IL-10) in the supernatants were measured using ELISAs (Invitrogen; Thermo Fisher Scientific) and gene expression was measured using RT-qPCR. The RT-qPCR protocol was the same as that described in earlier publications (33–36) using the primers listed in Table I. Additionally, the tumoricidal activity of macrophages was evaluated using carboxyfluorescein diacetate succinimidyl ester (CFDA-SE)-labeled cancer cells, according to a previously established procedure (14). CFDA-SE (20 μM) in PBS was used to treat 1×10^5^ HCT116 cells for 30 min at 37°C, according to the manufacturer’s protocol. Passively diffused CFDA-SE in the cytoplasm was cleaved by intracellular esterase, resulting in fluorescence activity. CFDA-SE-labeled HCT116 cells (1×10^5^ cells/well) were then incubated with TAMs (1×10^5^ cells/well) with and without extracts (PL or PSS at IC_50_ values) for 72 h before being fixed with 2% paraformaldehyde and the nuclei stained with 4′, 6-diamidino-2-phenylindole (DAPI; Thermo Fisher Scientific). The fluorescence intensity was then calculated using ImageJ [National Institutes of Health (NIH), Maryland, USA].

### Animal and tumor xenograft model

The study protocol was approved by the Faculty of Medicine’s Institutional Animal Care and Use Committee at Chulalongkorn University (Bangkok, Thailand; ASP SST 018/2563), which followed the NIH in the United States. Male nude BALB/cAJcl-nu mice aged 8 weeks were obtained from Nomura Siam International Co., Ltd. (Bangkok, Thailand). The mice were housed at a specific pathogen-free mouse facility in a controlled environment, with a temperature of 24±2°C, 50% relative humidity and a 12-h light-dark cycle, with light provided from 7:00 a.m. to 7:00 p.m. Additionally, 5 mice were housed in cages with corncob bedding, which was autoclaved before use. The mice had continuous access to a standard sterile diet and were provided autoclaved tap water from a bottle placed within the cage. To assess the *in vivo* effects of PL and PSS, a human colorectal cancer cell line (HCT116) was used as a representative tumor. HCT116 cells (1×10^5^) in 100 μl DMEM supplemented with 10% FBS were subcutaneously injected into the mice, with or without daily intralesional injections of extracts (PL or PSS at 2 mg in 200 μl) or normal saline (control group), which started 2 weeks after cancer inoculation. Total of 15 mice were used for subcutaneous cancer injection, with 5 mice for control and 5 mice per group for PL and PSS. The human endpoints include i) any one of severe tumor characteristics (higher than 4,000 mm^3^ of tumor volume, ulcerated tumor, interference with eating or ambulation, and metastasis), ii) weight loss more than 20%, and iii) moribund appearance (labored breathing, loss of righting reflex, lethargy, ascites, and severe diarrhea). However, no mice reached the human endpoint, and all mice were observed for 30 days. For anesthesia during the subcutaneous injection of tumor cells, 3% isoflurane was adjusted to 0.5-0.75% v/v with an oxygen-nitrous oxide mixture (VetFlo™ Isoflurane Vaporizers) (Kent Scientific, Torrington, CT, USA) during the injection procedure. Notably, 3% isoflurane was released into the induction box to initiate the anesthetic process (anesthesia induction) and maintain by diluting isoflurane into 0.5-0.75% using the machine during the injection process (anesthesia maintenance). During the short anesthesia for cancer injection, the mice were placed on a temperature control bed (37 °C) and checked for the depth of anesthesia (righting and palpebral reflexes, muscular tone, and respiratory rate) before the injection. The tumor volume (mm^3^) was determined using the following equation: (length x width x width)/2 (37). The control groups received subcutaneous injection of the same volume of autoclaved dH_2_O. Tumor measurement, as well as routine health examinations of the mice, including assessment of body weight, was performed every 3 days. All of the mice were sacrificed through cardiac puncture under isoflurane anesthesia (the induction and maintenance process) at 3 weeks after intralesional administration. Moribund mice, as indicated by the presentation of severe pain, severe distress or suffering conditions, including infection, chronic inflammation, loss of mobility, abnormal resting posture, losing >20% of their weight or a tumor diameter of >2 cm, were considered humane endpoints, after which the mice were sacrificed by cardiac puncture under isoflurane anesthesia.

*Statistical analysis.* All data were subjected to statistical analysis using the Statistical Package for Social Sciences software (SPSS 22.0; SPSS Inc., Chicago, IL, USA) and Graph Pad Prism version 10.0 software (Dotmatics, La Jolla, CA, USA). The results are presented as the mean ± standard error. Statistical significance between groups was assessed using unpaired t-test for two-group comparisons, or one-way analysis of variance (ANOVA) with Tukey’s test for multiple-group comparisons. The time-point experiments were analyzed using repeated measures ANOVA. P<0.05 was considered to indicate a statistically significant difference.

## Results and discussions

### Impacts of PL and PSS extracts against cancer cell lines

The effects of PSS and PL extracts were compared to cisplatin on colorectal cancer (HCT116, SW620, and HT-29), prostate cancer (PC3), and kidney (HEK 293) cell lines, particularly regarding cytotoxicity, cell cycle arrest, and cell death. The MTT assay was used to assess cytotoxicity by measuring cell viability following treatment with the extracts and cisplatin in a dose-dependent manner, to ascertain the most suitable conditions for subsequent investigations. Table II shows the estimated IC_50_ values for the extracts and cisplatin against all cell lines, as calculated using a linear equation of inhibitory concentrations. As a result, cisplatin revealed cytotoxicity against all tumor cell lines and kidney cell lines at the lowest concentration, followed by PSS, whereas PL showed cytotoxicity only at high dosages (Fig. 1A-E). In terms of PSS, the results showed that it was the most dangerous to SW620 cells, and the least detrimental to HEK 293 cells (Fig. 1E and Table 2).

**Figure 1.**
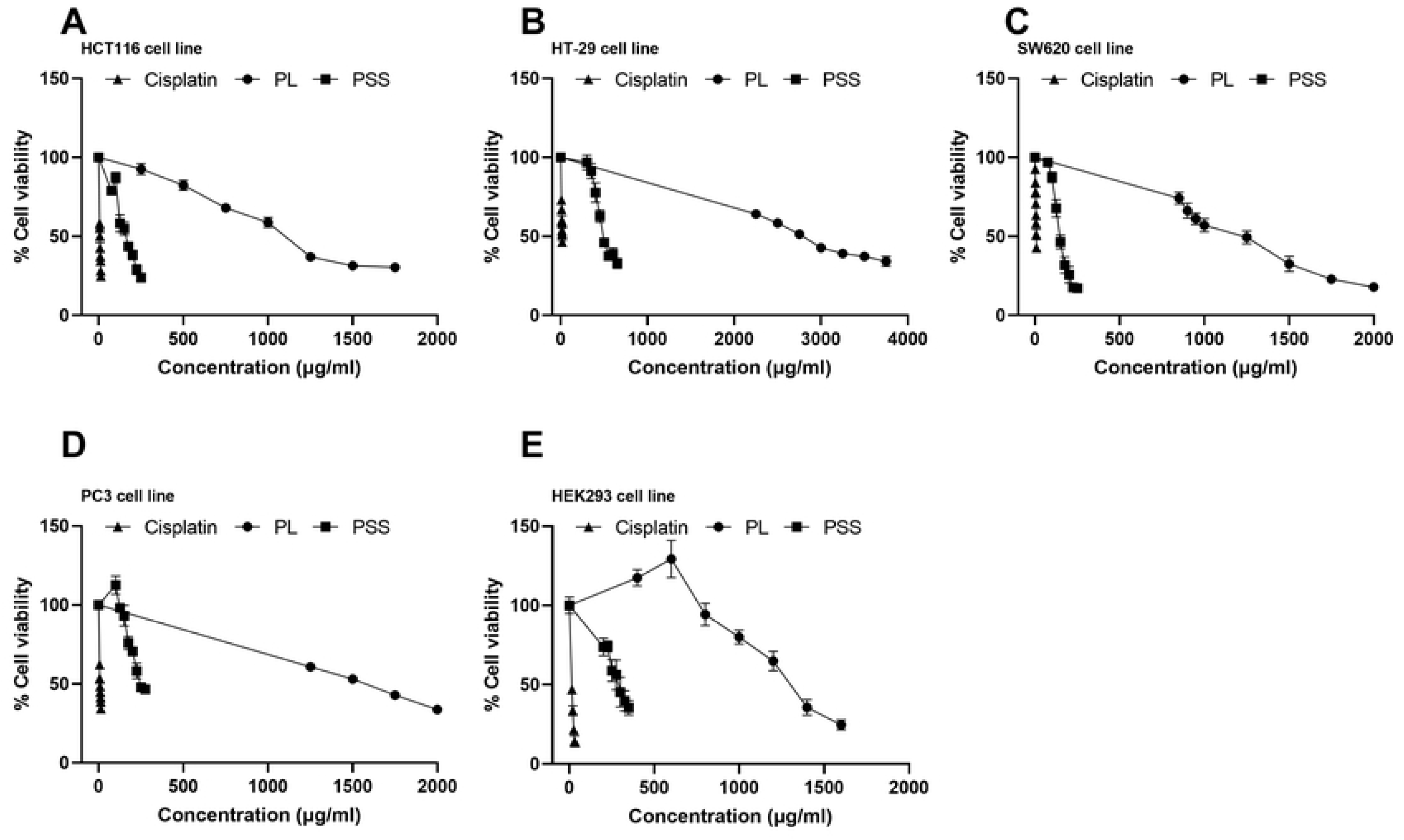
Characteristics of the impacts of compounds from PL and PSS mixture and Cisplatin against colon cancer cell line including HCT116 (A), HT-29 (B), SW620 (C) prostate cancer cell line PC3 (D) and PSS against kidney cell line HEK293 (E) as indicated by the dose-dependent cytotoxicity. *, p < 0.05 between the indicated groups; #, p < 0.05 vs. the untreated.

**Table 2.** The 50% inhibitory concentration (IC_50_) of the tested agents against several cell lines.

| Cancer cell lines | IC<sub>50</sub> (95% CI), R square |  |  |
| --- | --- | --- | --- |
|  | PL extract (ug/mL) | PSS extract (ug/mL) | Cisplatin (ug/mL) |
| HCT116 | 1,087 (1,025 – 1,132), 0.9 | 159 (150 – 168), 0.8 | 6.6 (6.2 – 7.0), 0.8 |
| HT-29 | 2,804 (2,721 – 2,884), 0.8 | 517 (499 – 537), 0.9 | 15.7 (14.2 – 18.1), 0.7 |
| SW620 | 1,163 (1,112 – 1,216), 0.8 | 150 (144 – 156), 0.9 | 7.2 (6.7 – 7.8), 0.9 |
| PC3 | 1,532 (1,472 – 1,586), 0.9 | 259 (238 – 265), 0.8 | 7.4 (7.0 – 7.7), 0.8 |
| HEK | 1,302 (1,207-1,409), 0.8 | 288.9 (270.6-312.7), 0.6 | 8.3 (6.4 – 10.4), 0.95 |

In the present study, three colorectal and one prostate cancer cell lines were tested, despite the availability of numerous cancer cell lines for research purposes. Among the three colorectal cancer cell lines, HCT116 is a highly aggressive cell line without the capacity to differentiate into enterocytes, whereas SW620 cells show resistance to glutamine-based targeting treatment, and HT-29 has an intermediate capacity to differentiate into enterocytes, is sensitive to glutamine-based therapy, and demonstrates AMP-activated protein kinase activation (38, 39). Hence, these colorectal cancer cell lines represent different characteristics of colon cancer. In parallel, the most commonly used cell lines for prostate cancer research are derived from different metastatic sites; for example, lymph nodes (LNCaP cells), vertebrae (PC3), and brain (DU145) (40). Because androgen receptor-positive prostate cancer can be partially treated with hormone therapies, novel additional therapies may be useful for the treatment of androgen receptor-negative cancer (PC3 and DU145) (41). Subsequently, PC3 was selected as a representative androgen receptor-negative prostate cancer cell line in this study. The impacts of herbal extracts, including PL and PSS, were assessed on both cancer cell lines and mice, as a proof of concept of using the *in vitro* test to select the use of adjuvant cancer therapy *in vivo*. Both PL and PSS are Thai remedies used for the treatment of several diseases, which might have value in modern medicine, as previously mentioned in breast cancer (12). While SC and SG are climber plants, PL is a mushroom that predominantly contains polysaccharides, especially β-glucan, with anti-proliferation and immune modulation properties (14). However, PL not only contains β-glucan, but also consists of several phenolic compounds that possibly synergistically attenuate cancer (42), as indicated by our earlier investigation that showed LC/QTOF-MS chromatographic peaks (10). As such, most of the possible active compounds in PL and PSS are similar. In PL and PSS, there were several substances with previously mentioned anticancer properties; for example, iridin, ethyl caffeate, sparteine, matrine, formononetin, rhoifolin, rosmanol and angelol (12). Only four compounds, including ethylcamptothecin, octadecanoic acid, methocyindole acetic acid, and brevicarine have been found in PSS but not in PL, representing the substances from SC and SG in the PSS mixture. Camptothecin is a quinolone alkaloid that has been used as a chemotherapeutic agent in the treatment of leukemia (43). Meanwhile, fewer mentions have been made regarding the anticancer effects of octadecanoic acid (18-carbon chain saturated fatty acid, which is referred to as stearic acid) and indole-3-acetic acid (the most common plant hormone of the auxin class that regulates various aspects of plant growth and development) (44, 45). In the present study, the PSS and PL extracts exhibited cytotoxicity against colorectal and prostate cancer cells. Extracts are harmful to various cancer cell lines. This finding is consistent with the research conducted by Florento et al., which discovered that there were varying concentrations of anticancer drugs in cancer cell lines (46). Cancer cells have different genetic characteristics from normal cells, and they also vary among cancer cell types. Various cell lines have diverse histological and genetic properties, which result in a response to varying extract doses. Many studies have demonstrated that the drug sensitivity of cancer cells is dependent on alterations to the genome, such as point mutations, copy number changes, and epigenetics (47, 48). Herbal medications have diverse anti-cancer effects on different cells as a result of mutations in aberrant signaling cells that might contribute to cancer. In this study, HEK293 represents normal kidney cells. Although HEK 293 is not a good representation of normal cells because it is an immortal human embryonic kidney cell, it is not cancerous. The findings suggested that PSS was more deleterious to HEK293 than HT29. It is possible to conclude that PSS is not appropriate for the treatment of HT29. Taken together, it is crucial to recognize that this herbal medicine is not suitable for all individuals if it is to be used in the clinic.

To evaluate cancer cell cycle arrest following treatment with IC_50_ doses of PSS, PL, and cisplatin, PI staining and flow cytometry were used to determine the DNA content of each cell line. Cell cycle arrest is a halting point in the cell cycle that signifies that the cell is unable to continue dividing. The results revealed that cisplatin caused an accumulation of all cancer cells in the G_2_ phase, as well as a decrease in the accumulation of all cancer cell lines in the G_0_/G_1_ and S phases when compared to the untreated control group. Meanwhile, PSS induced G_2_ cell cycle arrest in all colorectal cancer cell lines, but not in PC3 prostate cancer cells, whereas PL induced G_2_ cell cycle arrest in just HT-29 and SW620 cells. All agents (cisplatin, PSS, and PL) resulted in no accumulation of cells in the G_0_/G_1_ and S phases (Fig. 2). Cisplatin was shown to be the most efficient therapy in producing early apoptosis in all cancer cell lines (10-20%), whereas PL and PSS resulted in ∼5% HT-29 cell apoptosis. Meanwhile, PL, but not PSS, resulted in 1-2% early PC3 cell apoptosis. Regarding the late apoptosis in all cancer cell lines, with cisplatin being the most powerful inducer in HCT116 and SW620 cells. Cisplatin and PL both triggered late apoptosis in HT-29 cells at equal rates. Only cisplatin and PL induced late apoptosis in the PC3 cell line, with cisplatin being more effective than PL. Across all groups, <5% of HCT116 cells treated with cisplatin displayed necrosis (Fig. 3). As a result, PL and PSS had less cytotoxic action on cancer cells than cisplatin. Following exploration of the cell cycle, the present study showed that cisplatin was the most effective agent in inducing cancer cell toxicity through G_2_ arrest, followed by PSS and PL, respectively. Notably, all of the tested agents did not induce the accumulation of cells in G_0_/G_1_ and S phases, indicating that G_2_ phase interference may be an underlying mechanism. Because of the well-known association between cell cycle arrest and apoptosis (49), it is not surprising that cisplatin strongly induced cell death (early apoptosis, late apoptosis, and necrosis). The present study demonstrated that cisplatin was more effective in inhibiting cancer cell proliferation than PSS and PL. PSS could induce apoptosis in cancer cells at a low rate (<5%), indicating that the potential of the extracted samples to induce apoptosis in these cancer cell lines was low. This study found that PSS was not an effective therapy for androgen receptor-negative cancer (PC3). Due to the study’s limitations, only PC3 represented this kind of prostate cancer. Other androgen receptor-negative cancer cell lines are offered as subjects to determine the efficacy of PSS.

**Figure 2.**
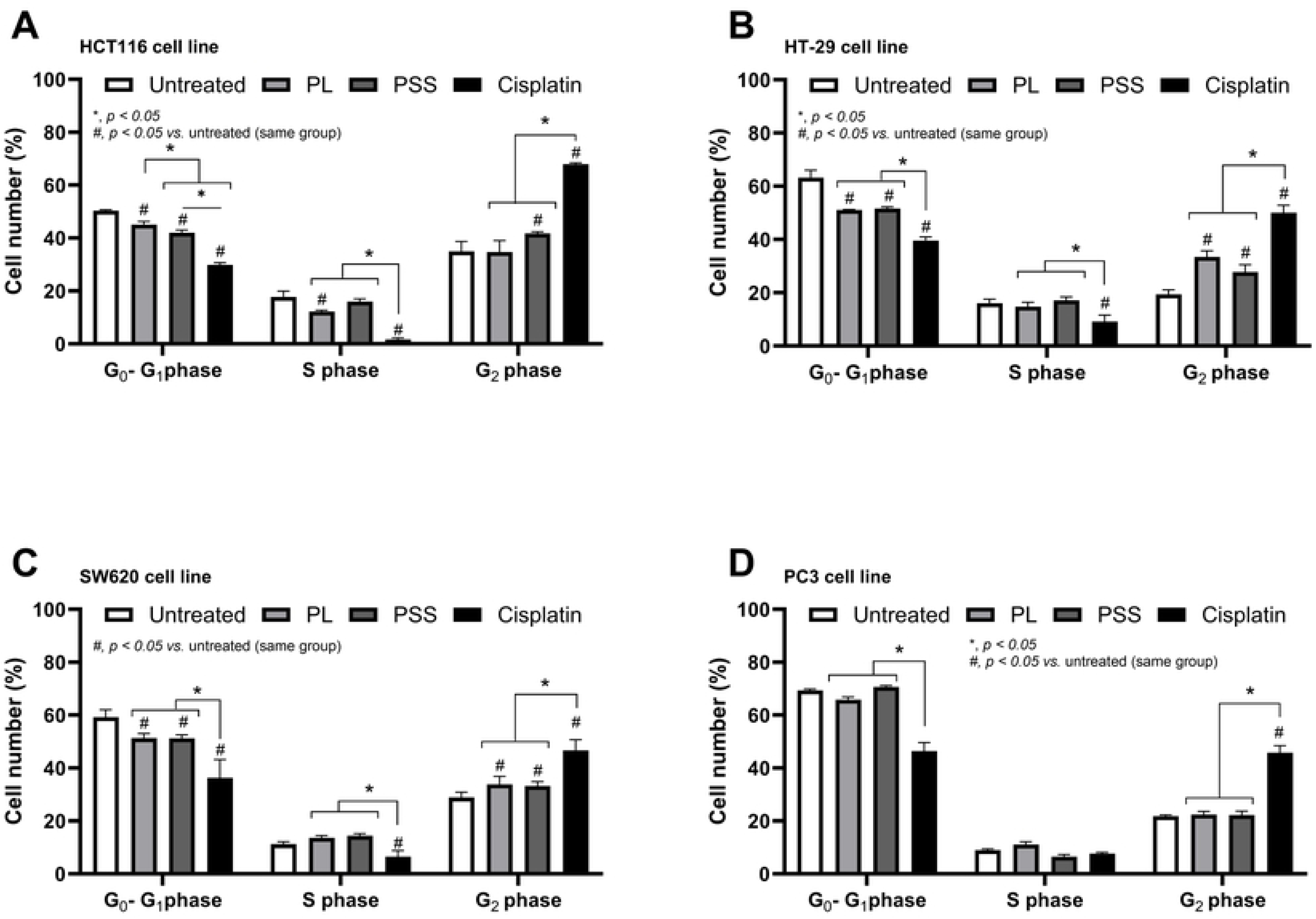
Characteristics of the impact of compounds from PL and PSS mixture (PL+ Smilax corbularia + Smilax glabra at the ratio 3:1:1) and Cisplatin against colon cancer cell lines including HCT116 (A), HT-29 (B), and SW620 (C) prostate cancer cell line PC3 (D) as indicated by cell cycle arrest. *, p < 0.05 between the indicated groups; #, p < 0.05 vs. the untreated

**Figure 3.**
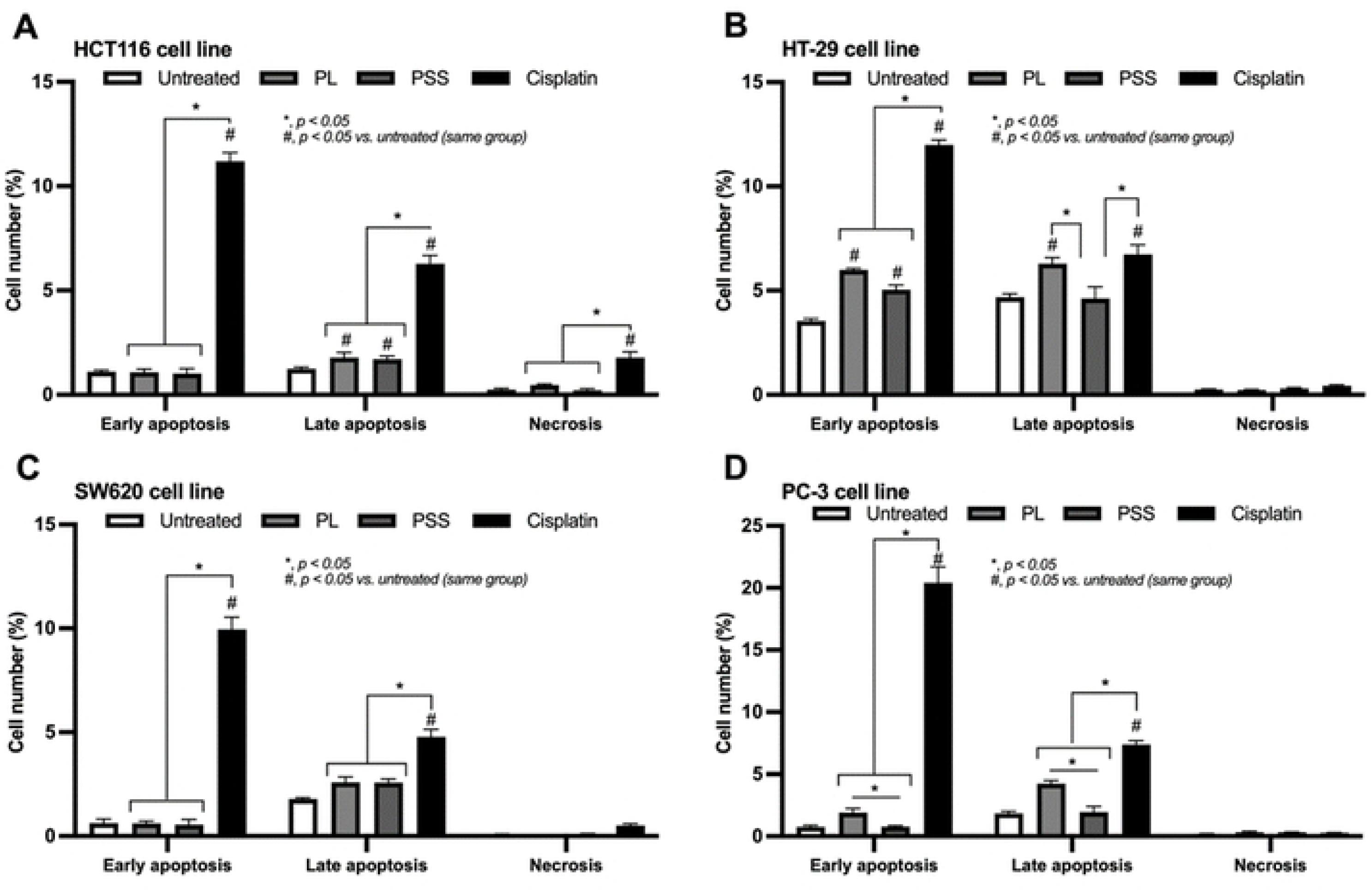
Characteristics of the impact of compounds from PL and PSS mixture (PL+ Smilax corbularia + Smilax glabra at the ratio 3:1:1) and Cisplatin (a standard cancer treatment) against colon cancer cell lines including HCT116 (A), HT-29 (B), and SW620 (C) prostate cancer cell line PC3 (D) as indicated by cell death. *, p < 0.05 between the indicated groups; #, p < 0.05 vs. the untreated

Unlike cisplatin, which has well-known cytotoxicity mechanisms through nuclear DNA binding and subsequent interference with cell transcription and DNA replication (50), the mechanisms of PL and PSS against malignant cells are still unknown and require further exploration. Regarding the underlying mechanisms, the induction of cell cycle arrest and cell death by PSS may be due to the impact of some compounds on the downregulation of several genes, including *MDM2*, *KRAS,* and *MKI67*. Due to the low potency of the anticancer effect of PL compared with PSS, the anticancer compounds of interest might be one or a combination of the four compounds that are found in PSS but not in PL. While detailed mechanistic exploration was not the primary aim of the present study, the impacts of the active compounds might partly be associated with these three selectively explored genes in the cells that may be involved in the cytotoxicity of PSS. Notably, *MDM2* upregulation has been mentioned in several malignancies, and has been reported to help cancer escape p53 surveillance, leading to the immortalization of the cells (51). Furthermore, *KRAS* and *MKI67* may increase cell proliferation, and the overexpression of these genes has been shown to enhance the growth rate of cancer (52, 53).

Accordingly, to explore the possible mechanisms of PL and PSS against these cancer cells, the expression of several genes associated with cancer cell death and cell proliferation, including *MDM2*, *KRAS*, and *MKI67,* was measured by RT-qPCR. The results demonstrated that both PL and PSS reduced *MDM2* expression across all cancer cell types, as well as *MKI67* expression in HCT116, HT29, and PC3, and *KRAS* expression in HT-29 and PC3. Only PSS decreased *MKI 67* expression, but PL marginally enhanced it in SW620. Unexpectedly, both PL and PSS boosted *KRAS* expression in HCT116. In SW620, only PL increased *KRAS*, but PSS had no effect on its expression (Fig. 4). Our findings revealed that PSS was more toxic in HCT116 (IC_50_=159 ug/mL) and SW620 (IC_50_=150 ug/mL) than in HT29 (IC_50_=517 ug/mL) and PC3 (IC_50_=259 ug/mL). However, at PSS’s cytotoxicity effect on each cell line did not correlate with a molecular level that detected the expression of *MDM2*, *KRAS*, and *MKI67*. *MDM2* expression was the most promising gene associated with cytotoxicity. *MDM2* expression in cells treated with PSS, compared to untreated cells, was reduced in HCT116 and SW620. Considering cytotoxicity, PSS exhibited toxicity toward HCT116 and SW620. This may suggest a possible correlation between the cytotoxicity of PSS and the downregulation of *MDM2* expression in both cells. In contrast, *MDM2* expression in PC3 following PSS treatment was reduced less than in HT29, whereas PSS is more cytotoxic in PC3 than in HT29. *KRAS* expression was the most unexpected finding. Its expression was lowered in HT29 and PC3 following PSS treatment compared to untreated, however, it increased in HCT116 and remained unchanged in SW620 after PSS treatment, in contrast to PSS’s cytotoxicity against these cell lines. One potential explanation is that HCT116 and SW620 contain the *KRAS* mutation, whereas HT29 and PC3 do not. Consequently, the herbal medicine does not have an impact on *KRAS* mutation cells. *MKI67* expression was lower in all cancer cell lines following PSS treatment compared to untreated. Its expression was the strongest decrease in HT29 cells, contradicting PSS’s cytotoxicity towards the cells, since PSS had the lowest cytotoxicity against HT29. The findings differed from our prior breast cancer study, which found reduced expression of *MDM2*, *KRAS*, and *MKI67* in breast cancer cell lines treated with PSS compared to untreated cells (12). These data suggest that PSS affects distinct cells with diverse genetic backgrounds. Taken together, the findings indicate that the molecular-level effect of PSS in this study is questionable; just three genes were discovered, which may be inadequate. To identify the PSS-affected genes, the transcriptome should be examined.

**Figure 4.**
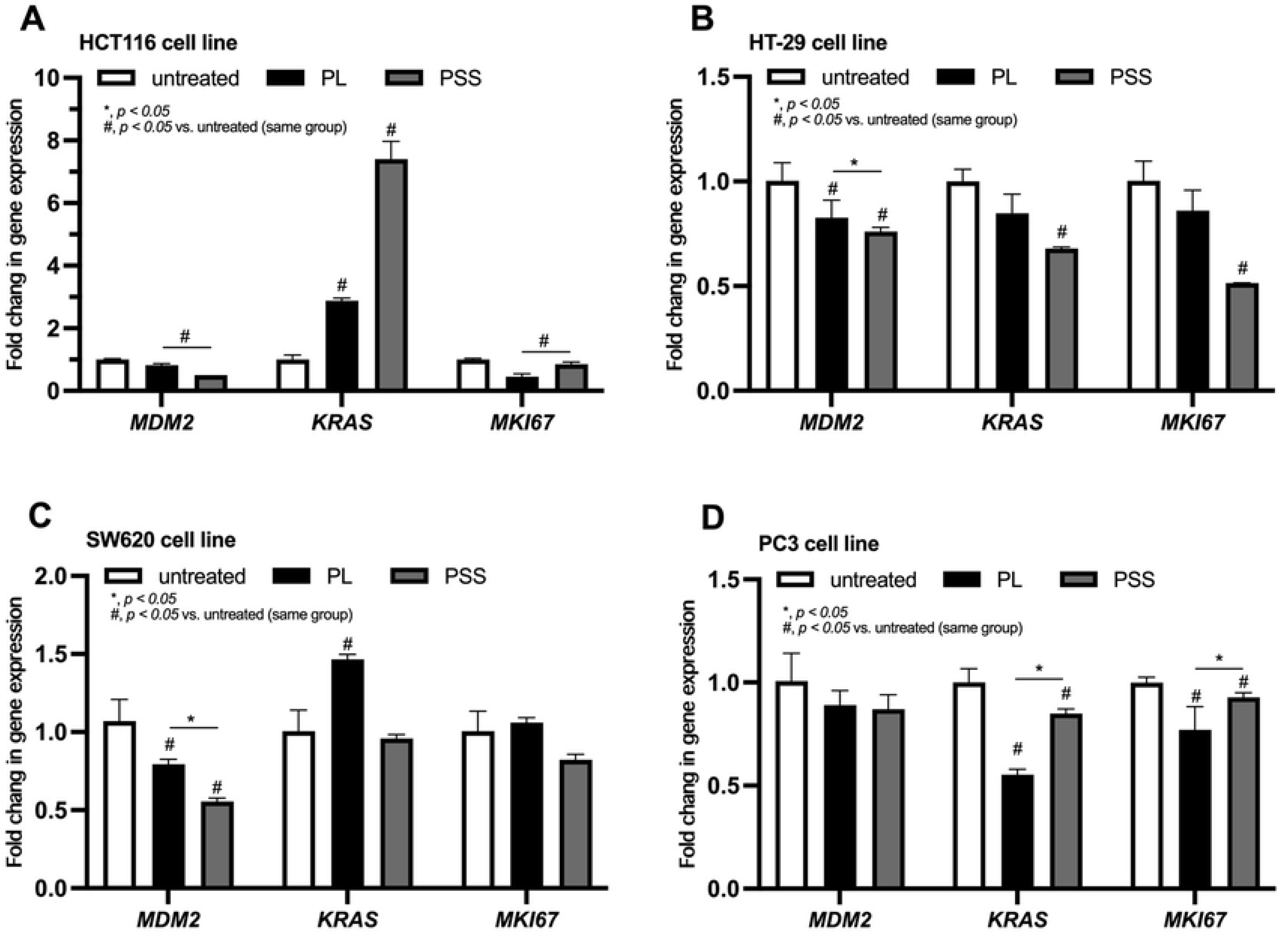
Characteristics of the impacts of compounds from PL and PSS mixture (PL+ Smilax corbularia + Smilax glabra at the ratio 3:1:1) and untreated control (untreated) against colon cancer cell lines, including HCT116 (A), HT-29 (B), and SW620 (C) and prostate cancer cells (PC3) (D) as indicated by the expression of several genes, including murine double mi-nute 2 (MDM2), Kirsten rat sarcoma virus (KRAS), and marker of proliferation Ki-67 (MKI67), of the malignant cells are demonstrated. Data derived from isolated triplicate experiments. *, p < 0.05 between the indicated groups; #, p < 0.05 vs. the untreated (within the same gene)

*Influences of PL and PSS extracts on TAMs.* The impacts of the extracts on cancer are possibly not only caused by cell cytotoxicity but may also be due to the regulation of TAMs, which are immune cells facilitating tumor growth. In the present study, supernatants from a colorectal cancer cell line (HCT116) were used to induce TAMs, whereas macrophages in control culture medium were used as naïve macrophages (M0). Regarding cytokines, TAMs upregulated IL-6 and IL-10 levels, and downregulated TNF-α when compared with M0 macrophages (Fig. 5A-C), supporting the role of IL-6 and IL-10 elevation, and TNF-α downregulation in cancer promotion (54, 55). Treatment with both extracts downregulated the levels of all of these cytokines (TNF-α, IL-6 and IL-10) (Fig 5A-C), which might have an anticancer effect (reduced IL-6 and IL-10) or could induce cancer promotion (decreased TNF-α). Regarding macrophage polarization, treatment with the cancer cell line supernatant downregulated genes associated with M1 pro-inflammatory polarization (IL-1β and iNOS) and upregulated genes associated with M2 polarization (Fizz-1, TGF-β and Arg-1) (Fig. 5F-H), thus supporting the M2-like condition of TAMs (56). In comparison with TAMs, both extracts altered the characteristics of TAMs, as indicated by elevated IL-1β and iNOS levels, and reduced Fizz-1 and TGF-β expression (Fig. 5D-G). However, only PSS, but not PL, downregulated Arg-1 expression (Fig. 5H) and effectively reduced the abundance of cancer cells after co-incubation with CSFE-stained cancer cells, as determined by the ratio of CFSE/DAPI fluorescence intensity (Fig. 5I and J). Notably, incubation of CSFE-stained cancer cells with M0 macrophages showed no difference compared with TAMs (Fig. 5I and J), despite the theoretical enhancing effect of TAMs on cancer growth (57). The anticancer effect is not only dependent on direct cancer cytotoxicity, but may also be due to an impact on immunity. As such, the effects of both PSS and PL on TAMs, the M2-like polarization of macrophages that can enhance tumor growth, and cytokine responses were also tested. Because TNF-α from macrophages might induce death processes in malignant cells (58), the downregulation of TNF-α in naive M0 macrophages in response to cancer cell conditioned medium might enhance tumor growth, thus supporting the role of TAMs in cancer (18). Likewise, the upregulated expression of *IL-6* and *IL-10* genes in TAMs might be beneficial for cancer, as IL-6 and IL-10 promote cancer cell proliferation and inhibit cancer cell apoptosis (55, 59). Notably, PL and PSS downregulated the expression levels of *IL-6* and *IL-10*, which might have a beneficial anticancer effect; however, the extracts also demonstrated a possible role in cancer promotion through the downregulation of TNF-α. On the other hand, the HCT116 colon cancer supernatant induced M2-like characteristics, as indicated by the upregulation of several genes associated with M2 polarization (*Fizz-1*, *TGF-β,* and *Arg-1*) and the reduced characteristics of M1 polarization (*IL-1β* and *iNOS* levels). Meanwhile, both PL and PSS similarly downregulated these M2 polarization-associated genes and upregulated the M1 polarization-associated genes (*IL-1β* and *iNOS*), which might be beneficial in patients with numerous types of cancer (57). Hence, PL and PSS demonstrated several antitumor effects, as indicated by downregulation of *IL-6* and *IL-10*, and the reduction in the direction of M2 polarization; however, they may also have had a tumor-promoting impact through the downregulation of TNF-α. Nevertheless, as determined in the CFSE-stained tumor cell experiments, only PSS, not PL, demonstrated an antitumor effect and reduced the abundance of the tumor cells, implying that the downregulation of TNF-α had a negligible effect on tumor promotion in this situation. Notably, PL did not exert an antitumor effect in the CFSE-stained tumor cell experiments, despite the similar impacts of PL and PSS on TAMs. Since Arg-1 expression was higher in TAMs after activation by PL compared with PSS, the reduced Arg-1 expression might be responsible for the greater antitumor effect of PSS over PL. Notably, the incubation of TAMs with the CFSE-stained tumor cells did not demonstrate any beneficial effects on the cancer cells, different from the well-known benefits of TAMs in cancer (18), suggesting a limitation of this experimental system. More mechanistic tests using other *in vitro* experiments may be interesting. The results of the *in vitro* macrophage experiments indicated that both PL and PSS might be candidates for additional anticancer therapies, and PSS demonstrated a more potent beneficial impact on some parameters. Hence, the different potencies between PSS and PL underscore the importance of doses and the intrinsic characteristics of the extracts.

**Figure 5.**
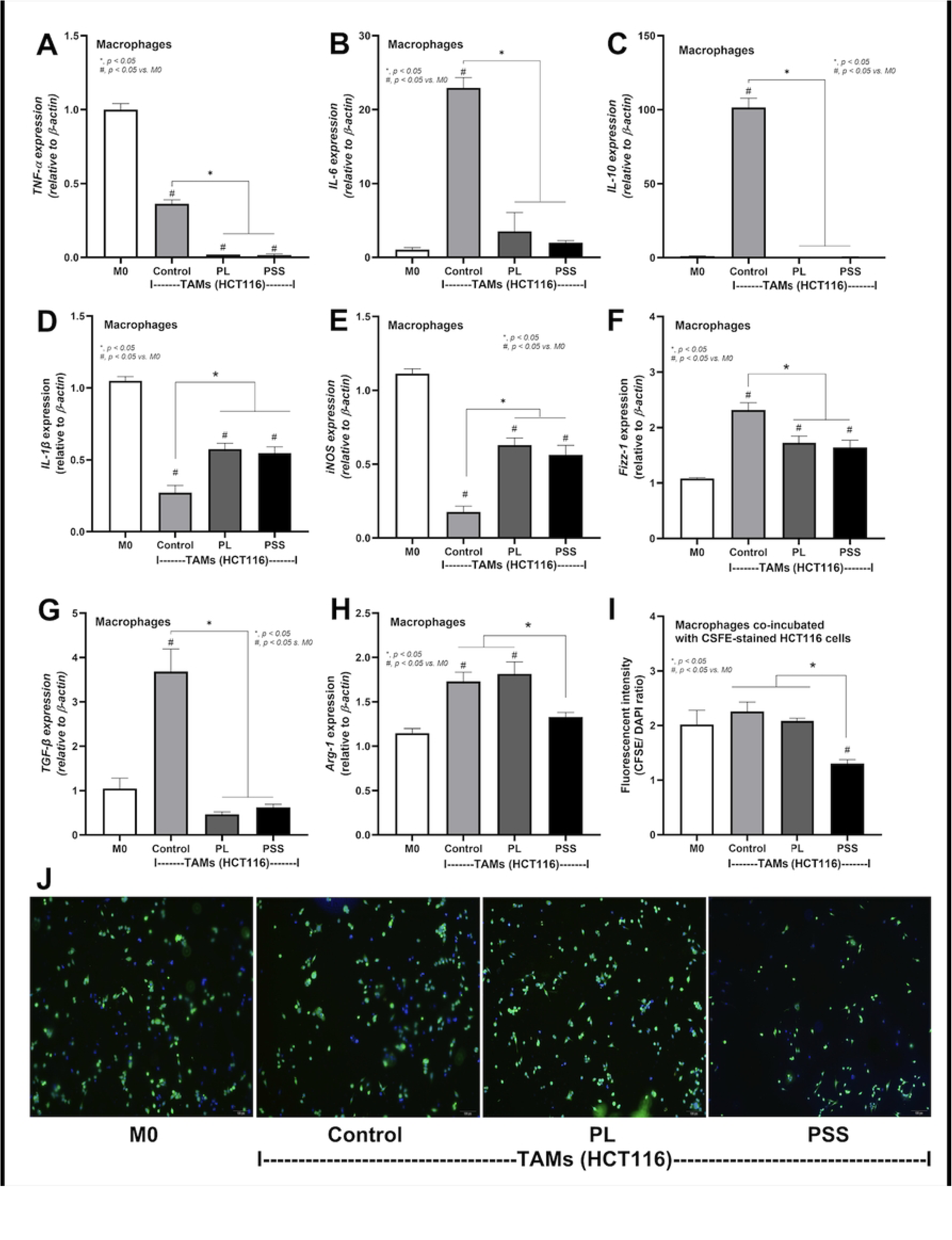
Characteristics of the impacts of compounds from PL and PSS mixture against TAMs using the supernatant from HCT116 as indicated by the expression of genes for cytokines (TNF-α, IL-6, and IL-10) (A-C), M1 macrophage polarization (IL-1β and iNOS) (D, E) and M2 macrophage polarization (Fizz-1, TGF-β and Arg-1) (F-H) with the cytotoxicity against carboxyfluorescein succinimidyl ester (CSFE)-stained malignant cells as indicated by fluorescent intensity and the representative pictures (I and J) are demonstrated. Data derived from isolated triplicate experiments. *, p < 0.05 between the indicated groups; #, p < 0.05 vs. M0

*Impacts of PL and PSS extracts on a mouse colon cancer (HCT116) xenograft model*. Because the extracts demonstrated some beneficial effects on cancer cell lines, especially colon cancer, the HCT116 colon cancer cell line was selected as a representative cell line for use in a mouse model of subcutaneous cancer injection (Fig. 6A). Notably, subcutaneous injection of HCT116 cells caused the generation of visible tumors in all mice, without alterations in body weight, as indicated in the positive control group (HCT116 cell injection with subcutaneous injection of sterile water) (Fig. 6B-E). Subsequently, the same dose of PL and PSS was administered in mice, with the maximum concentration that can be prepared into an injectable solution used (2 mg in 200 µl). Due to limitations of the solubility of the extracts, the prepared concentration of PL was 4-fold lower than the IC_50_ value of PL, whereas the concentration of the injectable PSS was equal to the IC_50_ value of PSS (Table II). Although there was no alteration in mouse body weights (Fig. 6B), the administration of PSS, but not PL, attenuated tumor volume, despite the fact that injection was performed 2 weeks after cancer injection (Fig. 6C-E). The higher IC_50_ value of PL, which may result in difficulties in the preparation of the extract in a soluble form, implies a possible limitation of the *in vivo* use of PL. In the present study, the HCT116 colon cancer cell line was selected as a representative cell line for the *in vivo* experiment, and only PSS, not PL, attenuated the tumors in mice. Indeed, the lack of anticancer properties of PL in mice might partly be due to the use of too low a concentration of PL *in vivo* due to the limited solubility of the high concentration of PL. Additionally, the benefits of PSS (PL + *Smilax* spp.) over PL alone might partly be due to the major compound of *Smilax* spp., especially camptothecin substances, which are not a component of PL. More studies on camptothecin against colon cancer may be interesting.

**Figure 6.**
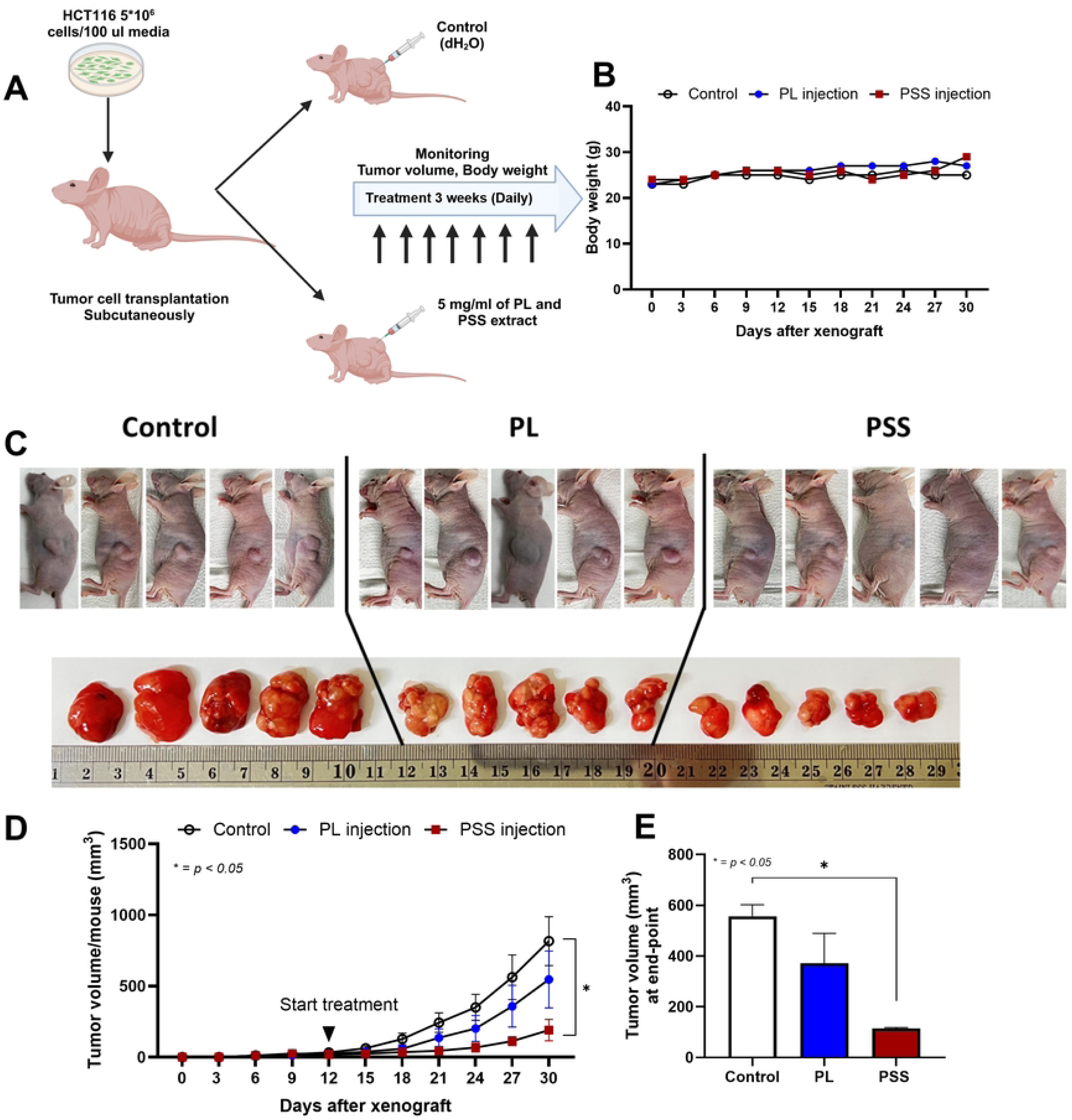
Characteristics of the impacts of compounds from PL or PSS mixture (PL+ Smilax corbularia + Smilax glabra at the ratio 3:1:1) and control (water injection) against sub-cutaneous colon cancer cell line (HCT116) as indicated by the schema of experiments (A), body weight (B), the tumor volume in time-point data and at the end-point (30 days post-cancer injection) with representative pictures of the tumors in mice and the pictures of tumors after excision (C-E) are demonstrated (n = 5 per time-point for B and D) (n = 5 per group for E). *, p < 0.05 between the indicated groups

Several limitations also need to be mentioned. First, different parts of the herbs might provide different bioactive agents (60), and the various solvents used in herbal preparation may be associated with the active ingredients (61). For example, the ethanol extracts of PL indicate strong antioxidant activity (62), whereas the aqueous PL extracts demonstrate more potent inhibitory effects on angiogenesis (63). The use of other solvents might demonstrate different effects on cancer. Second, the *in vivo* test (subcutaneous cancer cell injection) was performed only with HCT116 cells, and the combination of these extracts with standard chemotherapy was not tested. Also, the lack of animal experiments with PC3 cells is a limitation of the present study. Because of the metastatic characteristics of PC3 (a prostate cancer cell line) (64), the use of intralesional PL and PSS is not appropriate, and the development of proper drug delivery to different sites of metastasis is needed. More studies are required for the translation of the present findings into patient treatment. Third, the extracts were tested only in THP-1 cells (monocytes isolated from the peripheral blood of a patient with acute monocytic leukemia), which is a cancer cell line, not a normal cell line. In addition, the ratio of tumor cells to macrophages used was ∼1:3 followed our previous protocol (14); the results might be different with the different concentrations of the extract with various ratio between the malignant cells and TAM. Tests on primary normal noncancerous macrophages directly extracted from volunteers and the non-cancerous from organs (colons and prostate glands) might also have different results. Fourth, the mechanistic exploration was performed only regarding RNA expression, not at the protein level, and the extracts might affect cancer cells through direct cytotoxicity and indirect TAM inhibition. More studies aiming to explore the mechanisms, not only the proof of concept, will be interesting. Thus, the present results were only a proof of concept to use the PSS extract, adapted from a Thai remedy, in cancer. The concentrations of PSS and PL employed in the current study are lower than those that could be obtained through herbal therapies.

Administration of the extract through other routes, including oral and intravenous administration, a concentration higher than the IC_50_ value reported from this in vitro research is necessary. This is dependent on the pharmacokinetics of both extracts, which require more research to provide information for future therapy applications. Although the use of these extracts might not be ready to use as herbal oral therapy or tea-liked supplement, our proof-of-concept results supported a possible tumoricidal effect from these herbs derived from the Thai traditional medicine.

## Conclusion

In summary, the cytotoxicity of PSS and PL has been proven against a variety of cancer cell lines. Notably, in response to PSS, the inhibition of cell proliferation was more significant than the effects on cell apoptosis. The immune modulation effects on TAMs were also demonstrated, and the effects of PSS were more pronounced than those of PL. In addition, PSS inhibited the growth of colon malignancies in rodents as a proof of concept. Future assessment of the underlying mechanisms and the use of PSS and PL in other malignancies may be intriguing.

## Acknowledgements

The authors would like to thank the Center of Excellence in Molecular Genetics of Cancer and Human Disease (Chulalongkorn University, Thailand) for providing laboratory facilities.

This research is funded by Thailand Science Research and Innovation Fund Chulalongkorn University (207396), the Program Management Unit for Human Resources and Institutional Development, Research and Innovation (B16F640175 and B48G660112) with Rachadapisek Sompote Matching Fund (RA-MF-22/65, RA-MF-13/66, and RA-MF-eAsia), the National Research Council of Thailand (NRCT) (N41A640076), and the 90th Anniversary of Chulalongkorn University Fund (Rachadapisek Sompote Endowment Fund).

## Availability of data and materials

The datasets used and/or analyzed during the current study are available from the corresponding author on reasonable request.

## Authors’ contributions

CC conducted the experiments, analyzed the data and drafted the original manuscript. PV, PS, and PA conducted the experiments and analyzed the data. AL and PY designed the study, supervised the study, wrote the proposal for grants, and revised the manuscript. All authors have read and approved the final manuscript.

## Ethics approval and consent to participate

Not applicable.

## Patient consent for publication

Not applicable.

## Competing interests

The authors declare that they have no competing interests.

## Notes

### Competing Interest Statement

The authors have declared no competing interest.

